# Microfluidic Quaking-Induced Conversion (Micro-QuIC) for Rapid On-Site Amplification and Detection of Misfolded Proteins

**DOI:** 10.1101/2023.07.17.549283

**Authors:** Dong Jun Lee, Peter R. Christenson, Gage Rowden, Nathan C. Lindquist, Peter A. Larsen, Sang-Hyun Oh

## Abstract

Protein misfolding diseases, such as prion diseases, Alzheimer’s, and Parkinson’s, share a common molecular mechanism involving the misfolding and aggregation of specific proteins. There is an urgent need for point-of-care (POC) diagnostic technologies that can accurately detect these misfolded proteins, facilitating early diagnosis and intervention. Here, we introduce the Microfluidic Quaking Induced Conversion (Micro-QuIC), a novel acoustofluidic platform for the rapid and sensitive detection of protein misfolding diseases. We demonstrate the utility of our technology using chronic wasting disease (CWD) as a model system, as samples from wild white-tailed deer are readily accessible, and CWD shares similarities with human protein misfolding diseases. Acoustofluidic mixing enables homogeneous mixing of reagents in a high-Reynolds-number regime, significantly accelerating the turnaround time for CWD diagnosis. Our Micro-QuIC assay amplifies prions by an order of magnitude faster than the current gold standard, real-time quaking-induced conversion (RT-QuIC). Furthermore, we integrated Micro-QuIC with a gold nanoparticle-based, naked-eye detection method, which enables visual discrimination between CWD positive and negative samples without the need for a bulky fluorescence detection module. This integration creates a rapid, POC testing platform capable of detecting misfolded proteins associated with a variety of protein misfolding diseases.

**TOC graphic:** 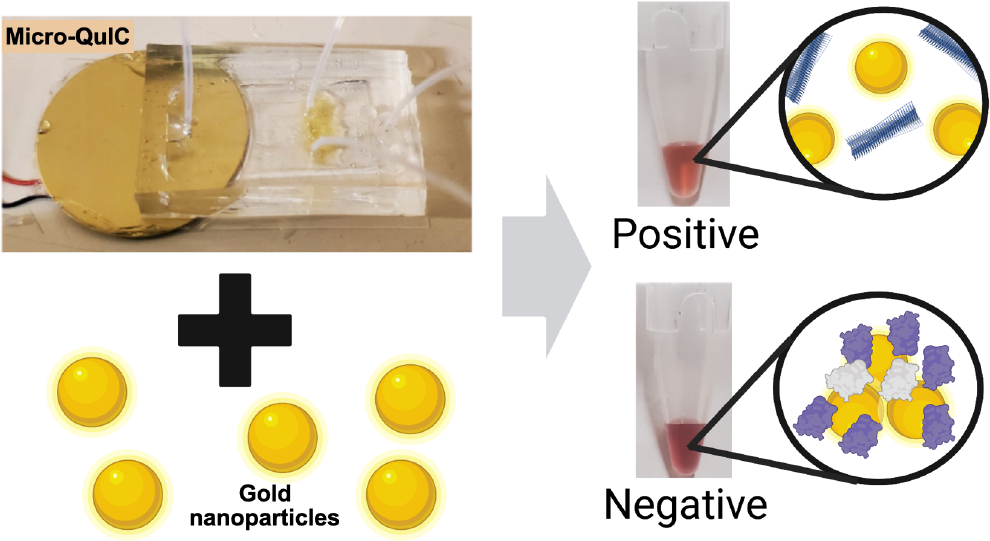

---

Protein misfolding diseases, including prion diseases (PDs), Alzheimer’s, and Parkinson’s, are a class of progressive neurodegenerative disorders that affect both animals and humans. The development of point-of-care diagnostic (POC) technologies for the accurate detection of misfolded proteins is essential for early diagnosis and intervention in these diseases. Prion diseases, a subset of protein misfolding diseases, are 100% fatal and impact both humans and animals, with examples including Creutzfeldt-Jakob disease (CJD) and fatal familial insomnia (FFI) in humans, bovine spongiform encephalopathy (BSE; also known as ‘mad cow disease’) in cattle, scrapie in sheep, and chronic wasting disease (CWD) in deer and elk^1-3^. These diseases are caused by the accumulation of infectious protease-resistant forms of the prion protein (PrP^Res^) within the central nervous system (CNS). A concerning feature of PrP^Res^ is that the infectious proteins withstand traditional methods of decontamination (e.g., standard autoclaving) and can remain infectious within a given environment for years^4-6^. Once formed, PrP^Res^ interacts with native cellular prion protein (PrP^C^), inducing misfolding events that ultimately lead to the creation of additional PrP^Res^ and related protein fibrils and plaques^7^. After onset, aggregated PrP^Res^ propagates throughout the CNS, killing neurons and supporting cells, and eventually cascading to death.

PDs are 100% fatal and typically have variable incubation times depending on the species impacted and underlying genetic factors^2^. The native cellular form of PrP^C^ is highly conserved across mammalian species, thus the potential exists for cross-transmission of PrP^Res^, the most notable example being the outbreak of variant CJD in humans after consumption of BSE contaminated meat products throughout the United Kingdom in the late 1980’s^8^. This research focuses on CWD as a model protein misfolding disease to develop a diagnostic technology, as CWD shares similarities with human protein misfolding diseases and samples from wild white-tailed deer are readily accessible. There is growing concern that CWD will cross multiple animal and human species barriers, leading to sporadic outbreaks of prion diseases in humans. With the global spread of CWD to cervid populations, including North America, Scandinavia, and South Korea, the number of exposure events to infectious CWD prions in both animals and humans is predicted to increase substantially^9^. Therefore, there is an urgent need for on-site diagnostic tools that can rapidly, sensitively, and accurately detect CWD-causing prions, ultimately benefiting the broader field of protein misfolding disease diagnostics.

Conventional diagnostic methods for CWD, such as antibody-based enzyme-linked immunosorbent assay (ELISA) and immunohistochemistry (IHC), are both time-consuming and expensive, and require substantial training and expertise to operate^10^. Moreover, their diagnostic sensitivity is limited due to the inability of commonly used antibodies to distinguish between PrP^C^ and PrP^Res^,^11^ thus necessitating complicated enzymatic, chemical, and/or heat digestion to enrich PrP^Res^. Therefore, a definitive diagnosis of CWD often requires post-mortem histopathological examination^12^.

A promising development in PD diagnostics involves the use of ultrasensitive seeding assays that facilitate the *in vitro* amplification of PrP^Res 13^. Real-time quaking-induced conversion (RT-QuIC)^14^ and protein misfolding cyclic amplification (PMCA)^15^ both exploit the ability of PrP^Res^ to induce PrP^C^ to misfold cyclically, forming quantifiable *in vitro* aggregates of protein fibrils. To achieve this, RT-QuIC utilizes shear force while PMCA utilizes sonication to mechanically fragment the PrP^Res^ fibrils into smaller nucleation sites. An important feature separating RT-QuIC from PMCA is that RT-QuIC conducts protein amplification using a recombinant prion protein substrate that does not produce infectious PrP^Res^ as the result of the assay. Alternatively, PMCA typically utilizes native hamster brain as a substrate, thus producing native misfolded PrP^Res^ throughout a given experiment. Another defining feature separating the assays is that RT-QuIC results are produced using standard fluorescent plate-readers that measure Thioflavin-T excitation levels with great sensitively throughout a given experiment, whereas traditional PMCA requires multiple rounds of Western-blotting and gel-image-based protein quantification. For these reasons, RT-QuIC is gaining more wide-spread usage for CWD research and diagnostics^16-19^.

While RT-QuIC boasts high sensitivity, its application for *in situ* or ‘on-the-spot’ CWD diagnosis remains curtailed by several factors. For example, it requires bulky and expensive equipment which can cost approximately $30,000-40,000 USD^20^. Moreover, manual handling of costly reagents increases the risk of contamination as well as reagent waste. Thus, there is a critical need to develop an *in situ* assay platform that confers rapid diagnosis of CWD in a fully automated manner while minimizing equipment size, cost and reagent usage.

In response to these challenges, microfluidics offers a promising avenue. Specifically, transitioning from macroscopic platforms to microfluidics can introduce numerous advantages for diagnostic application, such as lower reagent requirement^21^, increased specific surface-area-to-volume ratios^22^, biohazard containment^23^, and heightened heat and mass transfer rates^24^. While some microfluidic platforms have shown potential in reducing assay time^25^, they tend to suffer from an inherently low Reynolds number, which limits the mass transfer to a diffusion-limited regime^26^. Moreover, the use of elaborate pumps limits their application in on-the-spot diagnosis^27^.

Within the realm of microfluidics, both active and passive micromixers have been developed including, inertial^28^, acoustofluidic^29,30^, electrokinetic^31^ and magneto-hydrodynamics micromixers^32^. Among these, acoustofluidic micromixers have proven to be a powerful tool due to the high mixing index, low-cost of operation^33^, biocompatibility^34^, portability and the contact-free nature of the technology^35^. Specifically, lateral cavity acoustic transducers (LCATs) have been widely investigated^36-38^. Given these developments in acoustofluidic transduction mechanisms, we hypothesized that acoustofluidic microstreaming would generate sufficient shear stress to fragment non-covalent interaction between amyloid-enriched prion fibril subunits, laying the groundwork for a new platform for the rapid amplification of misfolded proteins.

Herein, we present an on-the-spot diagnostic platform integrating an assay similar to RT-QuIC, harnessing the power of acoustofluidic missing for PrP^Res^ and PrP^C^. Our device, shown in Fig. 1, comprises a PDMS-covered glass coverslip with reagent mixing facilitated by lateral cavity acoustic transducers, where arrays of dead-end side channels capture air bubbles which function as a vibrative membrane (Fig. 1b). Upon the application of a high frequency soundwave (4.6 kHz), the liquid-air interfaces resonate and the resulting acoustic field energizes the bulk liquid and produces a net force perpendicular to the bubble interface and out the end of the cavity^39^. The resulting acoustic streaming enhances the collision between PrP^Res^ and PrP^C^ as well as fragmentation of PrP^Res^ into smaller pieces, which in turn result in the multiplication of active seeds for the PrP^Res^ nucleation. LCATs are well suited for this application due to the simplicity in fabrication, tunability of mixing rate and high mixing index^35^. By combining the reagent microstreaming induced by the acoustofluidic micromixer and the quaking-based prion fibril amplification, we demonstrate that our method drastically reduces the amplification time from 48 hrs to 3 hrs. Furthermore, Micro-QuIC can be integrated with a gold-nanoparticle-based aggregation assay^20^, eliminating the need for bulky auxiliary detection modules. The Micro-QuIC device features advantages such as simplicity of use, automation, low cost, and portability, which are necessary for the future development of an automated all-in-one, on-chip amplification toolkit for *on-the-spot* diagnosis of protein misfolding diseases.

**Fig.1.**
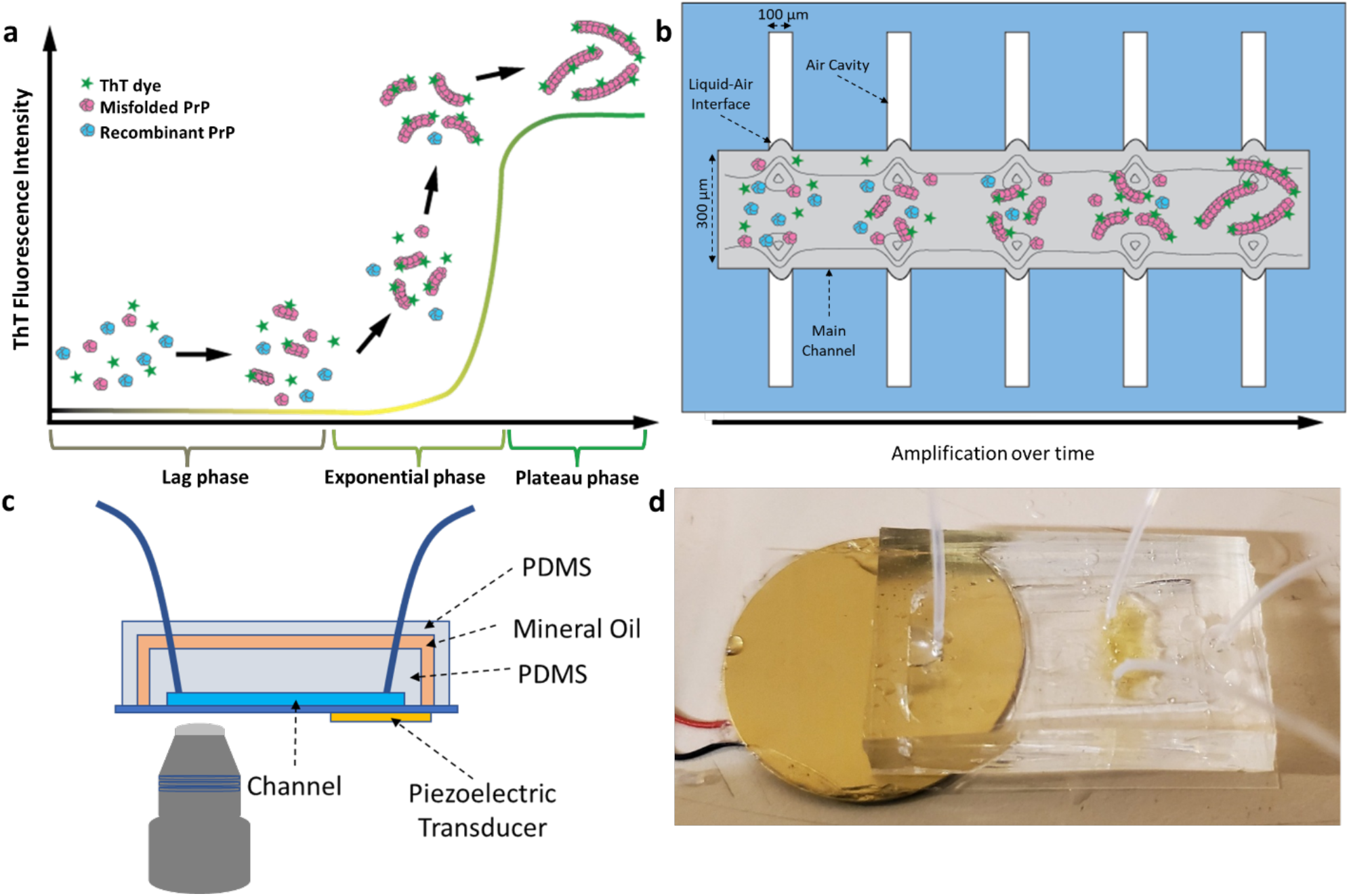
Overview of the Micro-QuIC system. A) Amplification of misfolded proteins. B) Misfolded chronic wasting disease (CWD) prion seeds originating from biological samples are added to rPrP solutions and injected into a microfluidic device. This creates a liquid-air interface between the main channels and side channels. The application of high frequency sound waves (4.56 kHz) induces vibration of the liquid-air interface to create a vortex current. These solutions are then shaken and incubated for approximately 3 hours at 45 ℃. If present, PrP^CWD^ induces conformational changes of the rPrP. Resulting products are labeled with Thioflavin T which can be detected under fluorescent microscope (excitation wavelength ∼ 450 nm, emission wavelength ∼ 427 nm.). C) Schematic of a Micro-QuIC device. Double layers of PDMS devices are bonded to the thin glass slides. Mineral oil is injected between the two layers of PDMS to prevent solution evaporation. A piezoelectric transducer is attached on the back of the glass slide. D) Photograph of a Micro-QuIC device.

## Results

### Microfluidic quaking induced conversion (Micro-QuIC) device

In this work, we sought to combine the active mixing benefits of acoustic technology with quaking induced prion amplification. To that end, Fig. 1c and d provide a schematic and photograph of the assembled Micro-QuIC device, respectively. The foundation of the acoustofluidic device is a thin glass coverslip that serves to transfer the vibrational energy from the acoustic transducer (shown in the figure) into the PDMS channels via flexural waves that travel out from the transducer and along the glass coverslip^40^. The primary design of the microfluidic channel geometry consists of lateral cavity structures, which capture air cavities upon injection of reagents. The resulting liquid-air interface serves as an oscillating membrane in response to vibration from the transducer, producing acoustic streaming in the channel to agitate the samples^39^.

The lateral cavity structure, with a channel width of 300 μm and a height of 100 μm, is repeated in 16 sets along the length of the main channel. Over the microfluidic PDMS layer, an outer PDMS channel creates a void filled with mineral oil, preventing heat-mediated evaporation of the reagent. The sample reagents are introduced into the main channel through the inlet while incubating the entire device at 45 **°**C. A piezoelectric transducer attached at the back of the glass slide generates an acoustic field to induce vibration of the liquid-air interface. This creates micro-vortex streaming in the main channel, which accelerates both the collision between PrP particles and the shear-induced fragmentation of PrP^Res^ fibrils. The collision between PrP particles in turn accelerates conversion of PrP^c^ to PrP^Res^, while the fragmented PrP^Res^ serves as a nucleation site for further fibril formation. As fibrils multiply, they bind to Thioflavin T, a fluorescent dye, rendering the amyloid aggregates visible under a fluorescent microscope.

### Device Characterization

Prior to prion amplification, we initially tested our device’s micro-vortex streaming capabilities using fluorescent microparticles. As shown in Fig. 2a and b, fluorescent microparticles showed vigorous circular motion upon application of an input bias of 10 V_pp_ at 4.6 kHz. The streaming velocity primarily depends on two parameters: frequency and voltage. To determine the frequency at which the LCAT generated the strongest acoustic streaming effect, the AC frequency was swept from 1 kHz to 100 kHz with a 50 Hz increment. Our experimental result indicated that the strongest acoustic streaming effect was generated when the device was excited at 4.6 kHz which coincided with the resonance frequency of the piezoelectric transducers. Outside the resonant frequency of the piezoelectric transducers, the streaming velocity drastically decayed (Fig 2c). Therefore, the AC bias was maintained at the resonant frequency for all subsequent experiments. The device was further characterized by applying different driving voltages to the piezoelectric transducer. Fig 2d shows the mixing performance with the different driving voltages at a frequency of 4.6 kHz. The results show that as the driving voltage of the piezoelectric transducer increased, the mixing efficiency was increased^39^.

**Fig.2.**
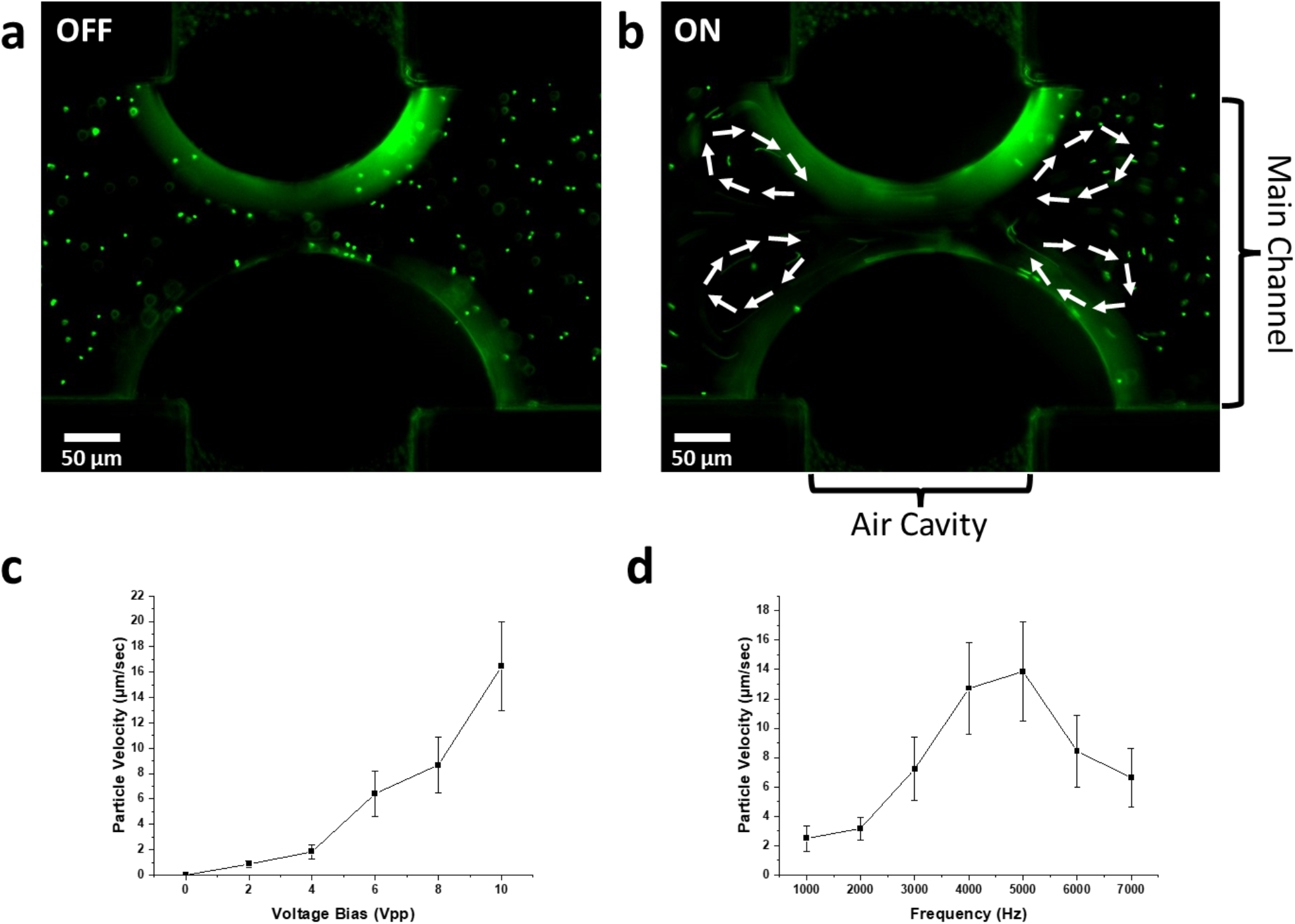
Actuation of microparticles. A) Fluorescent micrograph of microparticles when voltage bias was applied to the piezoelectric transducer. B) Fluorescent micrograph of microparticles with an input bias of 10 V_pp_ of 4.6 kHz. The trajectory of microparticle motion is illustrated in white arrows. C) Relationship between the applied voltage bias versus the average velocity of microparticles. D) Relationship between the applied frequency versus the average velocity of microparticles.

### On-chip Prion Amplification with Micro-QuIC

Next, we established the experimental condition for Micro-QuIC. Initially, a spontaneously misfolded rHaPrP (positive control) was spiked in a master mix consisting of 1X PBS, 1mM EDTA, 170mM NaCl, 10 μM ThT and rHaPrP ranging from 0.1 mg/mL to 0.4 mg/mL. We found that the use of a high concentration of the rHaPrP accelerates the reaction kinetics and hence the rHaPrP concentration was maintained at 0.4 mg/mL at all times unless stated otherwise. Both positively (misfolded) and negatively (non-misfolded) seeded master mixes were injected in a Micro-QuIC device and amplified for 180 min. Here, the 10 V_pp_ AC bias at a frequency of 4.6 kHz was applied for 30 min intervals with 30 min of rest. The fluorescent images were taken every 30 min to follow the amplification process over 180 min (Fig 3a). For the quantification of fluorescent intensity, we analyzed fluorescent particles in the field of view by thresholding. Both the number of fluorescent particles and their size increased from the positively seeded sample, indicating successful prion amplification, while there was no measurable change in fluorescence intensity in the negatively seeded sample). Based on our result, we have successfully demonstrated that prion amplification can be achieved in 180 min (Fig 3b) which is 16 times faster than conventional RT-QuIC (Fig 3c). To validate the amplified prion fibrils, we used transmission electron microscopy (TEM) to compare the dimensions and the morphology of amyloid fibrils amplified by both RT-QuIC and Micro-QuIC (Fig 3d and e). As reported previously^41^, the fibrils from both samples were helical in shape with an average width of 25 nm, which further validates that amplification products were equivalent in both methods.

**Fig.3.**
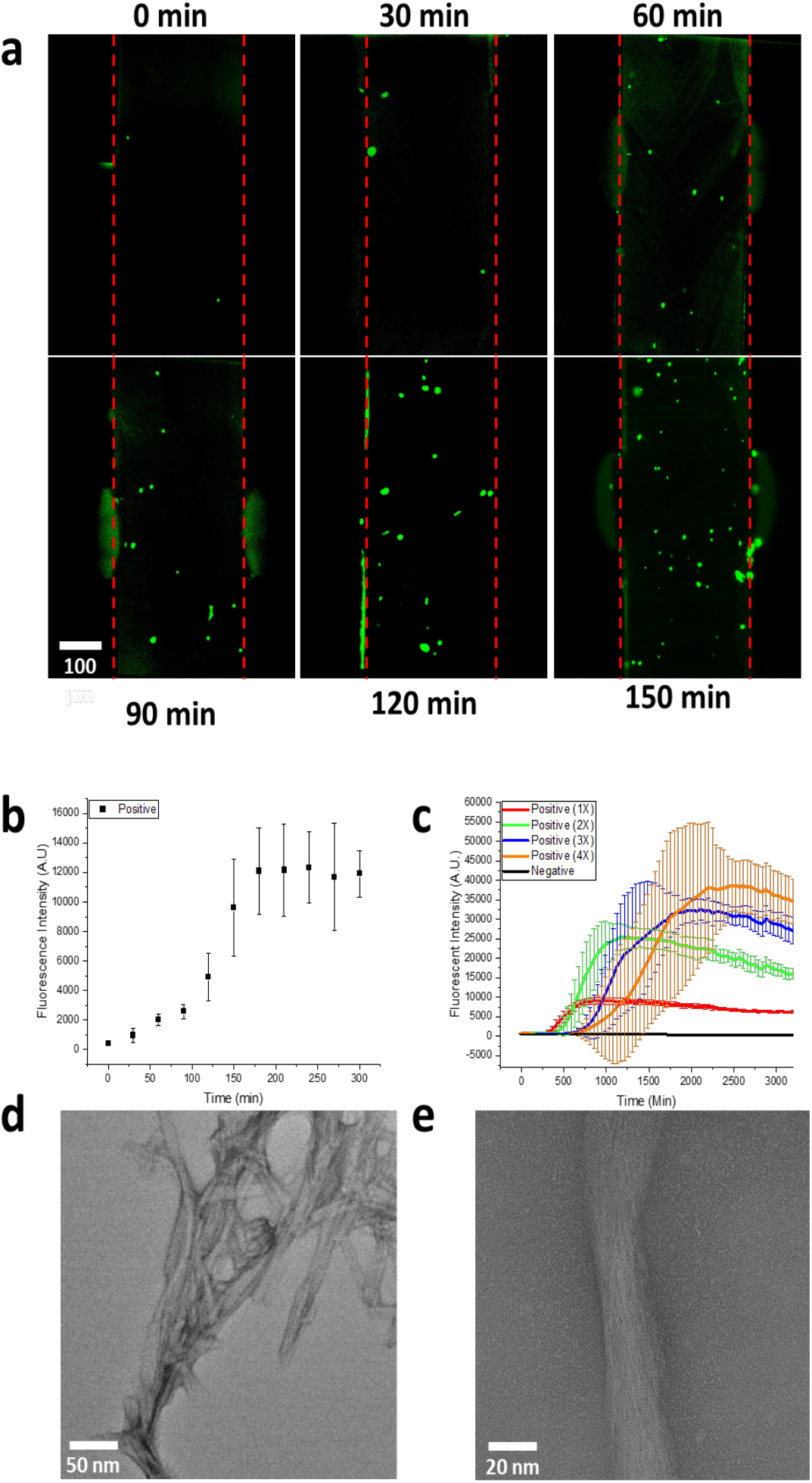
On-chip amplification of prion in real-time. A) Fluorescent micrograph of aggregated prion particles when periodic mixing with 30s interval was carried out by applying bias of 10 V_pp_ at 4.6 kHz from 0 min to 150 min. B) Fluorescence intensity of prion particles amplified by Micro-QuIC device over 300 min of amplification process. The fluorescence intensity was measured by particle tracking algorithm. C) Fluorescence intensity of prion particles amplified by conventional RT-QuIC place reader over 50 hrs. D and E) Morphology of prion fibrils amplified by Micro-QuIC device taken by TEM.

### On-site prion amplification and visual detection via Micro-QuIC and gold nanoparticle-based assay

Recognizing that Micro-QuIC can greatly accelerate rHaPrP misfolding and amplification kinetics, we explored its potential for field-portable diagnostics of misfolded proteins using PrP^CWD^ positive and negative white-tailed deer lymphoid tissues as model systems. We prepared homogenates of independently confirmed CWD positive and negative white-tailed deer medial retropharyngeal lymph nodes (RPLN) following methods as reported in Schwabenlander et al.^16^, then introduced them into Micro-QuIC devices along with the master mix. We tested positive (n=5) and negative (n=5) tissue samples. Each sample was incubated at 40 ℃ with periodic acoustic mixing as before for 3 hrs. End-point fluorescence measurement of the microfluidic channel showed that we were clearly able to distinguish CWD positive and negative samples through the difference in fluorescence intensity (Fig.4a). The significant reduction in assay time (∼3 hr) makes Micro-QuIC an attractive alternative to the current gold-standard for CWD diagnostics.

Christenson et al. previously reported the optical-based misfolded PrP detection system utilizing gold nanoparticles (AuNPs), which eliminates the need for a bulky and costly fluorescence detection module^20^. This system leverages the plasmonic properties of gold NPs^42,43,44^. Further, nanoparticles have been shown to influence amyloid formation kinetics^45,46,47,48,49^ and enhance the speed and sensitivity of conventional RT-QuIC^50^. In conjunction with Micro-QuIC, the AuNP based detection system would provide a valuable tool towards a field deployable platform for detecting misfolded proteins. To verify the compatibility of Micro-QuIC and AuNP assays, tissue samples were first amplified in a microfluidic channel at 40 ℃ with periodic acoustic mixing for 3 hrs as before. The post-amplified mixture was drawn from the microfluidic channel and incubated with a gold nanoparticle reagent for 10 min. Upon addition of positive samples, the absorbance peak remained unchanged at 515 nm while the addition of negative sample absorbance peaks was shifted to a longer wavelength of approximately 560 nm, which showed itself as a clear visual distinction between CWD positive and negative samples (Fig 4b). This result is consistent with previous results using the AuNP assay, thus demonstrating that our Micro-QuIC assay is effectively amplifying PrP^CWD^ associated with CWD positive samples and producing negative results for CWD negative samples. The integration of Micro-QuIC with nanoparticle-enhanced QuIC methods, such as Nano-QuIC or AuNP assay, suggests promising potential toward rapid, efficient amplification of misfolded proteins paired with a compact, visual detection method for point-of-care disease diagnostics.

**Fig.4.**
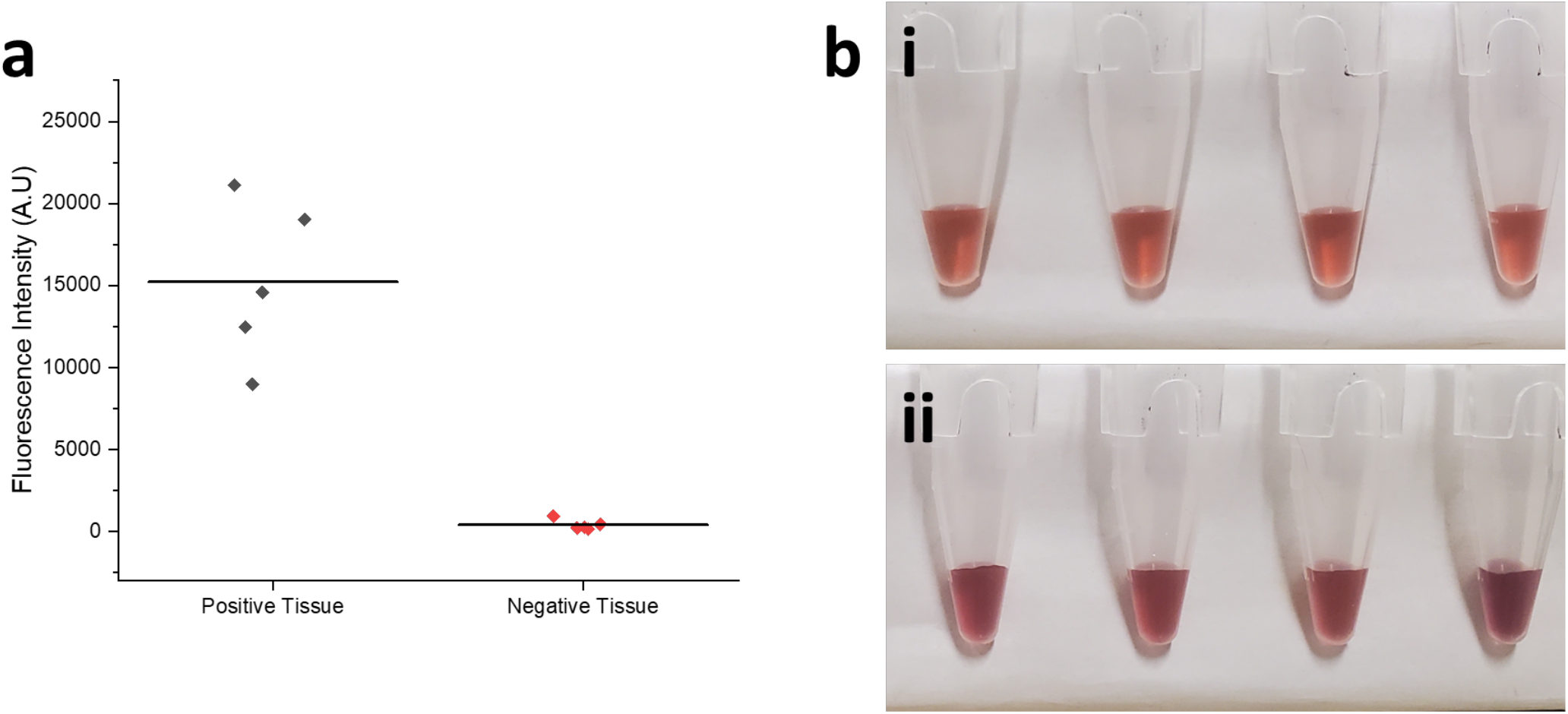
Amplification of real samples via Micro-QuIC and integration of AuNP-based visual detection. **a.** Measurement of fluorescent intensity of independently confirmed CWD-positive and CWD-negative white-tailed deer medial retropharyngeal lymph node (RPLN) tissue samples, amplified via the Micro-QuIC device. This figure depicts the significant differentiation in fluorescent intensity between the CWD positive and negative samples, providing a clear demonstration of the device’s amplification capabilities. Visual representation of optical-based misfolded PrP detection using gold nanoparticles. Gold nanoparticle (AuNP) based aggregation assay for the detection of amplified prion particles. (i) This panel shows that PrP^Res^, the pathogenic prion protein, does not interact with the surface of AuNPs and hence does not alter absorbance of AuNPs. (ii) Conversely, PrP^C^, the normal cellular prion protein, readily interact with the surface of AuNP to form aggregation, which cause blue shift of absorbance, making it visually distinct and detectable with naked eye.

## Discussion

The historic vCJD outbreak in the United Kingdom following BSE prion exposure underscores the necessity for routine and effective PD surveillance strategies. It is within this framework that concern is escalating across North America, Scandinavia, and South Korea as CWD continues to spread within cervid populations that are distributed across expansive geographic regions and that are a common source of food, food supplements, or herbal medicine. Recent estimates indicate that venison from tens of thousands of CWD positive cervids are unknowingly being consumed^9^, resulting in frequent exposure events to infectious prions and testing the human/cervid species barrier. For this reason, there is an urgent need to develop innovative diagnostic tools capable of the accurate, rapid, and ‘on-the-spot’ detection of CWD. Our approach combines RT-QuIC amplification of CWD prions with an acoustofluidic micromixer for swift amplification of target misfolded proteins. We hypothesized that an acoustofluidic micromixer would exert high shear stress to PrP^Sc^ fibrils, inducing active fragmentation, which in turn serve as new nucleation sites. Our strategy has successfully slashed the diagnosis time to approximately 3 hours, a significant improvement over standard RT-QuIC.

We have further enhanced the utility of our approach by integrating a AuNP-based visual detection method into our amplification strategy. This integration offers several benefits, notably making Micro-QuIC compatible with non-fluorescent-based detection, eliminating the need for bulky fluorescence detection modules, and enabling the rapid, portable, and *on-site* diagnosis of misfolded proteins. This opens up the potential for a field-deployable, POC testing platform, a development which could have far-reaching implications for protein misfolding disease diagnosis.

Our Micro-QuIC technology holds promise for the rapid, on-site, and visual identification of CWD positive and negative lymph tissues post-QuIC amplification. We chose RPLN and palatine tonsils collected from white-tailed deer for this study, as these tissues are ideal for early and accurate identification of CWD infection. Future analyses will focus on using Micro-QuIC for antemortem CWD diagnostics. RT-QuIC amplification protocols using samples acquired from living deer have recently been reported^18, 51, 52^ and these protocols could readily be combined with Micro-QuIC to provide rapid field-deployable ante-mortem tests of both wild and farmed cervids. Moreover, it is possible that Micro-QuIC has utility as a food-safety test given the recent documentation of RT-QuIC-based detection of CWD in white-tailed deer muscles that are used for human and animal consumption^53^. More broadly, we posit that our Micro-QuIC technology has the potential to be a versatile ‘on-site’ platform for detecting a variety of protein misfolding diseases where RT-QuIC and PMCA have been utilized, including scrapie in sheep, BSE in cattle, and CJD, amyotrophic lateral sclerosis, Parkinson’s, and Alzheimer’s in humans^54^.

## Notes

The authors declaring the following competing financial interest: a provisional patent with inventors, D.J. Lee, P.A. Larsen, S.-H. Oh, has been filed. Patent covers specific ideas, design, and protocols outlined within this paper. All other authors declare no competing interests.

## Acknowledgements

We thank Suzzanne Stone and Marc Schwabenlander for logistical assistance for the experiments reported herein. Manci Li and Marc Schwabenlander provided helpful comments. The Minnesota Department of Natural Resources kindly provided access to the white-tailed deer tissues used herein. Parts of this work were carried out in the Characterization Facility, University of Minnesota, which receives partial support from the NSF through the MRSEC (Award Number DMR-2011401) and the NNCI (Award Number ECCS-2025124) programs. Funding for research performed herein was provided by the Interdisciplinary Doctoral Fellowship from the University of Minnesota to D. J. L and P.R.C., the Minnesota State Legislature through the Minnesota Legislative-Citizen Commission on Minnesota Resources (LCCMR), Minnesota Agricultural Experiment Station Rapid Agricultural Response Fund (RARF), the Sanford P. Bordeau Chair in Electrical Engineering at the University of Minnesota to S.-H.O., and start-up funds awarded to P.A.L. through the Minnesota Agricultural, Research, Education, Extension and Technology Transfer (AGREETT) program.

## Supporting Information

The Supporting Information is available free of charge via the internet at http://pubs.acs.org

Methods section summarizing sample preparation and Micro-QuIC protocols.

