## Supplementary Information for "Microfluidic Quaking-Induced Conversion (Micro-QuIC) for Rapid On-Site Amplification and Detection of Misfolded Proteins"

#### Table of Contents

Experimental Methods
**Additional Statistical Information**
**Animal Research Statement**

### Methods

#### *Materials*

All Materials were used as purchased unless noted otherwise. SU8 2050 (Microchem Inc.) silicon wafer (Siegert Wafers), SU8 developer solutions (Microchem Inc.) microscope cover glass (24 X 50 mm, Globe Scientific Inc.) Piezoelectric transducers (7BB-27-4L0, Mouser electronics) Thioflavin T (ThT), 100 kDa Pall MWCO filter, Ethylenediaminetetraacetic acid (EDTA), sodium chloride (NaCl), phosphate buffered silane (PBS), trimethoxysilane were purchased from Sigma Milipore. Fluorescent microspheres (FCDG006) were purchased from Bangs Laboratories Inc. Polydimethylsiloxane (PDMS) and curing agent were obtained as SYLGARD®184 silicone elastomer kit from Dow Corning.

#### *Preparation of Recombinant Substrate*

The synthesis and purification of recombinant hamster PrP (HaPrP90-231) followed the methods of Schwabenlander et al. (2022). In brief, a truncated form (amino acids 90-231) of the Syrian hamster PRNP gene was cloned into the pD431-SR expression vector (ATUM, Newark, CA, USA) and was expressed in Rosetta (DE3) *E. coli* to synthesize the substrates. Subsequent purification and quality assessments followed Schwabenlander et al. (2022)[1].

#### *RT-QuIC for Spontaneous Misfolding of Recombinant Prion Protein*

For QuIC analysis, a master mix was prepared following specifications: 1X PBS, 1mM EDTA, 170mM NaCl, 10 µM thioflavin T (ThT), and 0.4 mg/mL rHaPrP. The 10% tissue homogenates were further diluted 100-fold in 0.1% SDS/PBS/N2 (final tissue dilution: 0.1%), and 2 µL of the diluent were added to each well containing 98 uL of RT-QuIC master mix. Spontaneous misfolding of recombinant prion protein was done similarly but with unfiltered recombinant proteins and reagents. For these reactions, no infectious seed was necessary because were run for <72 hrs. The spontaneously misfolded material was used to seed reactions for both Micro-QuIC and RT-QuIC. Plates were amplified on a FLUOstar® Omega plate reader

(BMG Labtech, Cary, North Carolina, USA) (42°C, 700 rpm, double orbital, shake for 57 s, rest for 83 s). Fluorescent readings were taken at ~45 min increments.

***Thermomixer-based Amplification***

We leveraged a standard benchtop shaking incubator (thermomixer) to produce QuIC-based prion amplifications as previously reported by Cheng et al.[2] and Vendramelli et al.[3], although with slight modifications. Plates which were made for amplification on the thermomixer were prepared identical to those amplified on the plate reader. Reactions were performed on a ThermoMixer® C equipped with SmartBlock plate and Thermotop (Eppendorf, Enfield, Connecticut, USA) at 48°C for 24hrs at 600 RPM (60s shake and 60s rest). We selected a 24 hour run time based on independent RT-QuIC results for lymph nodes and palatine tonsils from CWD+ white-tailed deer reported in Schwabenlander et al., including those examined herein, showing significant seeding activity within 9 to 24 hours (Fig. S3)[1].

***Preparation of Microfluidic Device and Operation***

The PDMS-based microfluidic device fabrication was performed using soft-lithography as described previously. Mater molds were prepared on 4-inch silicon wafers using a spin coating (Model CEE-100, Brewer Science Inc.) SU8 2050 to a height of 50 µm. Following the coating process, a pre-baking step was performed before UV-light exposure through a film mask with the requisite design, onto the SU8 coated silicon wafer for a duration of 12 s (MA6, Karl Suss). A post-baking step was performed before the SU8 development process. SU8 development was performed by gently washing the wafer in the developer solution for 3 min. Finally, the Si-wafer was silanized for 30 min with trimethoxysilane. PDMS-based microfluidic chips were produced by heat curing (90 °C for 3 h) PDMS with a curing agent mixture (10:1) on the master mold Si-wafer. Cured PDMS was peeled and cut into individual chips. Holes were punched at respective inlets and outlets using a 1 mm biopsy puncher (Acuderm Inc.) before bonding the PDMS to microscope cover glass. The bonding procedure was performed by treating both cover glass and the PDMS with a high-frequency generator (BD-10A, Electro-Technic Inc.) for 1 min. The microfluidic devices were heated for 2 h at 65 °C to help with the bonding process. For the fabrication of the outer PDMS casing, a master mold was fabricated by attaching a rectangular mold (3 cm x 1.5 cm x 0.5 cm) on a 4-inch silicon wafer, followed by silanization with trimethoxysilane for 30 min. Outer PDMS casing was fabricated by heat curing the PDMS master mix at 90 °C for 3 hr. Once cut into individual chip size, holes were punctured at respective inlets and outlets using a 1 mm biopsy puncher. Outer PDMS casing was bonded to a microfluidic device by aligning the inlets and outlets. Mineral oils were injected in between the microfluidic PDMS channel and the outer PDMS casing. Finally, piezoelectric transducers were attached at the back of the device using epoxy glue. The photograph of a final assembled device can be seen in Fig 1d. Before conducting the amplification experiments with recombinant PrP samples, all of our devices were experimentally tested with 0.005% fluorescent polystyrene microspheres (1 μm) to determine the optimized working frequencies of piezoelectric transducers to achieve uniform mixing. Based on these tests, 4.6 kHz was determined to be the working frequency for all our micro-QuIC devices; all the amplification experiments were conducted at this frequency. To track aggregated PrP, fluorescent images were obtained every 30 min for approximately 3 hr using laser excitation at 445 nm. The fluorescence intensity was measured by modifying the corrected total cell fluorescence measurement.

***Gold-Nanoparticle Based Detection***

Gold-nanoparticle-based prion detection followed the methods of Christenson et al.[4]. In brief, 2.45 nM 15 nm citrate capped gold nanoparticles (Nanopartz™, Loveland, Colorado, USA) were buffer exchanged to low concentration phosphate buffer (10mM $Na_2HPO_4$(Anhydrous), 2.7mM KCl, 1.8mM $KH_2PO_4$ (Monobasic)). After the amplification, protein solutions were diluted by two-fold in 1X PBS with the addition of final concentrations of 1mM EDTA, 170mM NaCl, 1.266mM Sodium Phosphate. Finally, 40ul of the dilute protein solution was then added to the 360ul AuNP solution and left to react at room temperature (RT) for 30 min. The color changes were observed both by naked eye and colorimeter (FLUOstar® Omega plate reader, BMG Labtech, Cary, North Carolina, USA) at 400-800 nm wavelength.

##### *Tissue Preparation*

Tissue preparation followed the methods of Christenson et al[4]. In brief, eight white-tailed deer tissues (4 CWD-negative and 4 CWD-positive) were obtained from white-tailed deer through collaboration with the Minnesota Department of Natural Resources and their CWD status was independently confirmed using the Bio-Rad TeSeE Short Assay Protocol (SAP) Combo Kit (BioRad Laboratories Inc., CA, USA) as well as RT-QuIC (Schwabenlander et al. 2022)[16]. White-tailed deer retropharyngeal lymph nodes (RPLNs) and palatine tonsils were homogenized in PBS (10 %w/v) with 1.5 mm zirconium beads with a BedBug Homogenizer (Benchmark Scientific, Sayreville, New Jersey, USA) on max speed for 90 s.

##### *Additional Statistical Information*

GraphPad Prism version 9.0 for Windows (GraphPad Software, San Diego, California USA, www.graphpad.com) was used for conducting statistical analysis. Three technical replicates were used to demonstrate the potential application of AuNP on spontaneously misfolded rHaPrP. Three and four technical replicates were used for testing CWD prions using AuNP and RT-QuIC, respectively for each animal. RPLN and/or palatine tonsil tissues from ten positive and fourteen negative animals were included in this study. The one-tailed Mann-Whitney unpaired u-test ($\alpha$=0.05) was used to test the average difference for all parameters of interests between samples.

##### *Animal Research Statement*

No deer were euthanized specifically for the research conducted herein and all tissues were secured from dead animals or loaned for our analyses. Thus, the research activities presented herein are exempt from review by the University of Minnesota Institutional Animal Care and Use Committee (as specified https://research.umn.edu/units/iacuc/submit-maintain-protocols/overview). White-tailed deer were euthanized for annual culling efforts to control the spread of CWD in Minnesota following Minnesota Department of Natural Resources state regulations and euthanasia guidelines established by the Animal

Care and Use Committee of the American Society of Mammalogists[5]. All methods and all experimental procedures carried out during this research followed University of Minnesota guidelines and regulations as approved by the Institutional Biosafety Committee under protocol #1912-37662H. This study was also carried out in compliance with the ARRIVE guidelines (https://arriveguidelines.org).
